# Selection on heterozygosity drives adaptation in intra- and interspecific hybrids

**DOI:** 10.1101/073007

**Authors:** Caiti S. Smukowski Heil, Christopher G. DeSevo, Dave A. Pai, Cheryl M. Tucker, Margaret L. Hoang, Maitreya J. Dunham

**Affiliations:** Department of Genome Sciences, University of Washington, Seattle, Washington, United States of America; Lewis-Sigler Institute for Integrative Genomics, Princeton University, Princeton, New Jersey, United States of America; Howard Hughes Medical Institute and Department of Embryology, Carnegie Institution, Baltimore, Maryland, United States of America; Department of Biology, Johns Hopkins University, Baltimore, Maryland, United States of America

**Author notes:** Current address: Exelixis, Dallas, Texas, United States of America (Christopher G. DeSevo). Current address: BioNano Genomics, San Diego, California, United States of America (Dave A. Pai). Current address: Wall High School, Wall Township, New Jersey, United States of America (Cheryl M. Tucker). Current address: Nanostring, Seattle, Washington, United States of America (Margaret L. Hoang). Corresponding author (MD).

## Abstract

Hybridization is often considered maladaptive, but sometimes hybrids can invade new ecological niches and adapt to novel or stressful environments better than their parents. However, the genomic changes that occur following hybridization and facilitate genome resolution and/or adaptation are not well understood. Here, we address these questions using experimental evolution of de novo interspecific hybrid yeast *Saccharomyces cerevisiae* x *Saccharomyces uvarum* and their parentals. We evolved these strains in nutrient limited conditions for hundreds of generations and sequenced the resulting cultures to identify genomic changes. Analysis of 16 hybrid clones and 16 parental clones identified numerous point mutations, copy number changes, and loss of heterozygosity events, including a number of nuclear-mitochondrial mutations and species biased amplification of nutrient transporters. We focused on a particularly interesting example, in which we saw repeated loss of heterozygosity at the high affinity phosphate transporter gene *PHO84* in both intra- and interspecific hybrids. Using allele replacement methods, we tested the fitness of different alleles in hybrid and *S. cerevisiae* strain backgrounds and found that the loss of heterozygosity is indeed the result of selection on one allele over the other in both *S. cerevisiae* and the hybrids. This illuminates an example where hybrid genome resolution is driven by positive selection on existing heterozygosity, and generally demonstrates that outcrossing need not be frequent to have long lasting impacts on adaptation.

## INTRODUCTION

Hybridization is now recognized as a common phenomenon across the tree of life. Historically however, the detection of hybrids has been difficult, and its incidence may be underreported for both plants and animals, and almost certainly for certain eukaryotes like insects and fungi (Albertin and Marullo 2012; Bullini 1994). Its importance as an evolutionary force has thus been maligned, as hybrids appeared both rare, and typically at a reduced fitness. In addition to potential post-reproductive barriers, the hybrid is theorized to be ill-adapted to its environment and will also suffer minority cytotype disadvantage because other hybrids are uncommon and backcrosses to parental species may be unfit (Mallet 2007). However, hybrids can have a variety of advantages over their parents, including heterozygote advantage, extreme phenotypic traits, and reproductive isolation (usually resulting from polyploidy), and can thus facilitate adaptation to novel or stressful conditions, invade unoccupied ecological niches, and even increase biodiversity.

Some hybridization events lead to new hybrid species (Mavarez, et al. 2006; Meyer, et al. 2006; Nolte, et al. 2005; Rieseberg 1997; Schumer, et al. 2014; Soltis and Soltis 2009), while most result in introgression from hybrid backcrosses to the more abundant parental species (Dasmahapatra, et al. 2012; Dowling, et al. 1989; Grant, et al. 2005; Taylor and Hebert 1993; Wayne 1993). This introduces genetic variation into a population at orders of magnitude greater than what mutation alone can achieve, in a sense operating as a multi-locus macro-mutation (Abbott, et al. 2013; Barton 2001; Grant and Grant 1994; Mallet 2007). Therefore, hybridization via introgression, polyploidy, or homopoloid hybrid speciation, may offer a rapid strategy for adaptation to changing environmental conditions. For example, in Darwin’s finches, adaptive introgression supplied the morphological variation which allowed the species to survive following an El Niño event (Grant and Grant 2010, 2002), while in ancient humans, introgression allowed adaptation to high altitudes (Huerta-Sanchez and Casey 2015), among other traits (Racimo, et al. 2015). The most iconic example comes from the hybrid sunflower species *Helianthus anomalus*, *Helianthus deserticola*, and *Helianthus paradoxus*, from the parents *Helianthus annuus* and *Helianthus petiolaris*. These three hybrid species are locally adapted to extreme desert, salt marsh, and dune habitats respectively, and show traits such as increased drought or salt tolerance relative to their parents (Heiser 1954; Rieseberg 1991; Rosenthal, et al. 2002; Schwarzbach, et al. 2001).

Agriculture and industry use both intra- and interspecific hybrids as a tool to increase yield or robustness, introduce resistance to pests, and create novel phenotype or flavor profiles. For example, plant breeders have crossed domesticated species to wild species to introduce resistance to a variety of pathogens in wheat, potato, and canola (Mason and Batley 2015), and almost all maize grown in the United States is grown from intraspecific hybrid seeds, which has increased yield and provided improved resistance to biotic and abiotic factors (Crow 1998). Vintners and brewers have created interspecific hybrids to select for traits such as lower acetic acid concentration (Bellon, et al. 2015), and many incidental fungal hybrids have been discovered in brewing and industry, including *Pichia sorbitophila* (Louis, et al. 2012b), and various hybrids across the *Saccharomyces* clade (Bellon, et al. 2015; Gonzalez, et al. 2006; Gonzalez, et al. 2008; Hittinger 2013; Muller and McCusker 2009), most notably the lager-brewing yeast, *Saccharomyces pastorianus* (Baker, et al. 2015; Dunn and Sherlock 2008; Gibson and Liti 2015; Peris, et al. 2016; Tamai, et al. 1998; Walther, et al. 2014). It is presumed that the severe selection pressures exerted during industrial processes have selected for interspecific hybrid genomes that may be more able to cope with the extreme environments.

At the genomic level, hybridization induces chromosome loss/aneuploidy, chromosomal rearrangements, gene loss, changes in gene expression, changes in epigenetic modifications, transposable element mobilization, and large scale loss of heterozygosity, in which the allele of one species is lost and the allele of the other species is retained via gene conversion or break induced replication (Abbott, et al. 2013; Ainouche and Jenczewski 2010; Albertin and Marullo 2012; Borneman, et al. 2014; Doyle, et al. 2008; Landry, et al. 2007; Masly, et al. 2006; Michalak 2009; Soltis, et al. 2014; Soltis 2013). These extensive changes can result in a chimeric, stabilized hybrid, although the period of time for genome stabilization to occur can range dramatically (Soltis, et al. 2014). There is some speculation that genetic distance between parental hybridizing species influences genome stabilization and bias in genome resolution, but this remains an open question. It is also unknown whether there are structural and functional biases in the ways in which genes/alleles are lost or modified. Both drift and selection influence the resolution of the hybrid genome, but their contributions are difficult to untangle.

To further explore these types of questions, an experimental approach is helpful. Researchers have long been exploring the genetics of hybrid traits in the lab, particularly in agricultural crops, although this is often slowed by infertility and reduced viability in many interspecific hybrids (Hajjar and Hodgkin 2007; Ouyang, et al. 2010; Perez-Prat and van Lookeren Campagne 2002). The genus of the budding yeast *Saccharomyces cerevisiae* lends itself particularly well to experimental study. Many hybrids of this genus have been discovered in brewing, industrial, and natural environments; indeed, the genus itself is speculated to be a product of an ancient hybridization event (Barbosa, et al. 2016; Hittinger 2013; Leducq, et al. 2016; Marcet-Houben and Gabaldon 2015). Viable interspecific hybrids can be created *de novo* in the lab (Greig, et al. 2002; Marinoni, et al. 1999), and their ability to grow mitotically means that the catastrophic postzygotic barriers to speciation that generally doom other obligate sexually reproducing hybrids can be avoided. This experimental system allows us to observe evolution in real time in the laboratory environment, and the genetic and genomic tools available in this model genus facilitate characterization of the connection between genotype and phenotype, including fitness.

Previous work in our lab group has utilized experimental evolution to investigate adaptive events in homozygous haploid and diploid *S. cerevisiae* (Gresham, et al. 2008; Payen, et al. 2014; Sunshine, et al. 2015). To investigate genome evolution post hybridization, we utilize an interspecific hybrid, *S. cerevisiae* x *Saccharomyces uvarum*, and its parentals: a homozygous diploid *S. uvarum* and an intraspecific hybrid S*. cerevisiae* GRF167 x *S. cerevisiae* S288C. This allows us to understand the impact of varying levels of heterozygosity on adaptation and genome evolution, ranging from none (*S. uvarum* and previous *S. cerevisiae* experiments), to intraspecific heterozygosity (S*. cerevisiae* GRF167 x *S. cerevisiae* S288C), to the most extreme case of interspecific hybrids. *S. uvarum* is one of the most distantly related species of *S. cerevisiae* in the *Saccharomyces* clade, separated by 20 my and 20% sequence divergence at coding sites (Cliften, et al. 2006; Kellis, et al. 2003). Despite this extensive divergence, *S. cerevisiae* and *S. uvarum* are largely syntenic and create hybrids, though less than 1% of zygotes are viable (Greig 2009). The two species differ in their stress tolerances, for example, *S. cerevisiae* being more thermotolerant, *S. uvarum* being cryotolerant (Almeida, et al. 2014). Previous evolution experiments using lab derived hybrids has revealed novel and/or transgressive phenotypes for ammonium limitation, ethanol tolerance, and growth on xylose (Belloch, et al. 2008; Dunn, et al. 2013; Piotrowski, et al. 2012; Wenger, et al. 2010). Notably, Dunn *et al.* (2013) reveal several loss of heterozygosity events and a repeatable reciprocal translocation that produces a gene fusion at the high-affinity ammonium permease *MEP2* after selection in ammonium limitation, offering insight into potential mutational events in the adaptation and/or stabilization of *S. cerevisiae* x *S. uvarum* hybrids.

Here, we evolved these hybrids and diploids in replicate in three nutrient limited conditions for hundreds of generations. Using whole genome sequencing, we found whole chromosome aneuploidy, genome rearrangements, copy number variants, *de novo* point mutations, and loss of heterozygosity. We sought to determine how initial heterozygosity impacts adaptation to novel conditions, and explore whether neutral or selective forces are influencing the resolution of the hybrid genome over time. In particular, we investigated a reoccurring loss of heterozygosity event observed in both intra- and interspecific hybrids, and found support for the hypothesis that loss of heterozygosity at this locus is due to selection.

## RESULTS

### Experimental evolution of hybrid and parental species

An interspecific hybrid was created by crossing *S. cerevisiae* and *S. uvarum* (strains in **Supplemental Table 1**), and evolved in continuous culture in the chemostat (Monod 1949; Novick and Szilard 1950a, b). In parallel, homozygous diploid *S. uvarum* and heterozygous diploid *S. cerevisiae* (GRF167xS288C) were also evolved. Each strain was grown in two or more replicate independent cultures under three different nutrient limitations—glucose, phosphate, and sulfate—for 85-557 generations (median 158) at 30°C, except for *S. uvarum*, which was unable to achieve steady state in all conditions at 30°C and so was evolved at 25°C. The population sizes were largely similar across strains, species, and conditions. Each evolved clone was subsequently competed individually against the appropriate GFP-tagged ancestor to gauge relative fitness. As expected, evolved hybrid and parental clones generally exhibit higher fitness than their unevolved ancestor, with typical relative fitness gains between 20-30% (**Tables 1**, **2**).

**Table 1:**
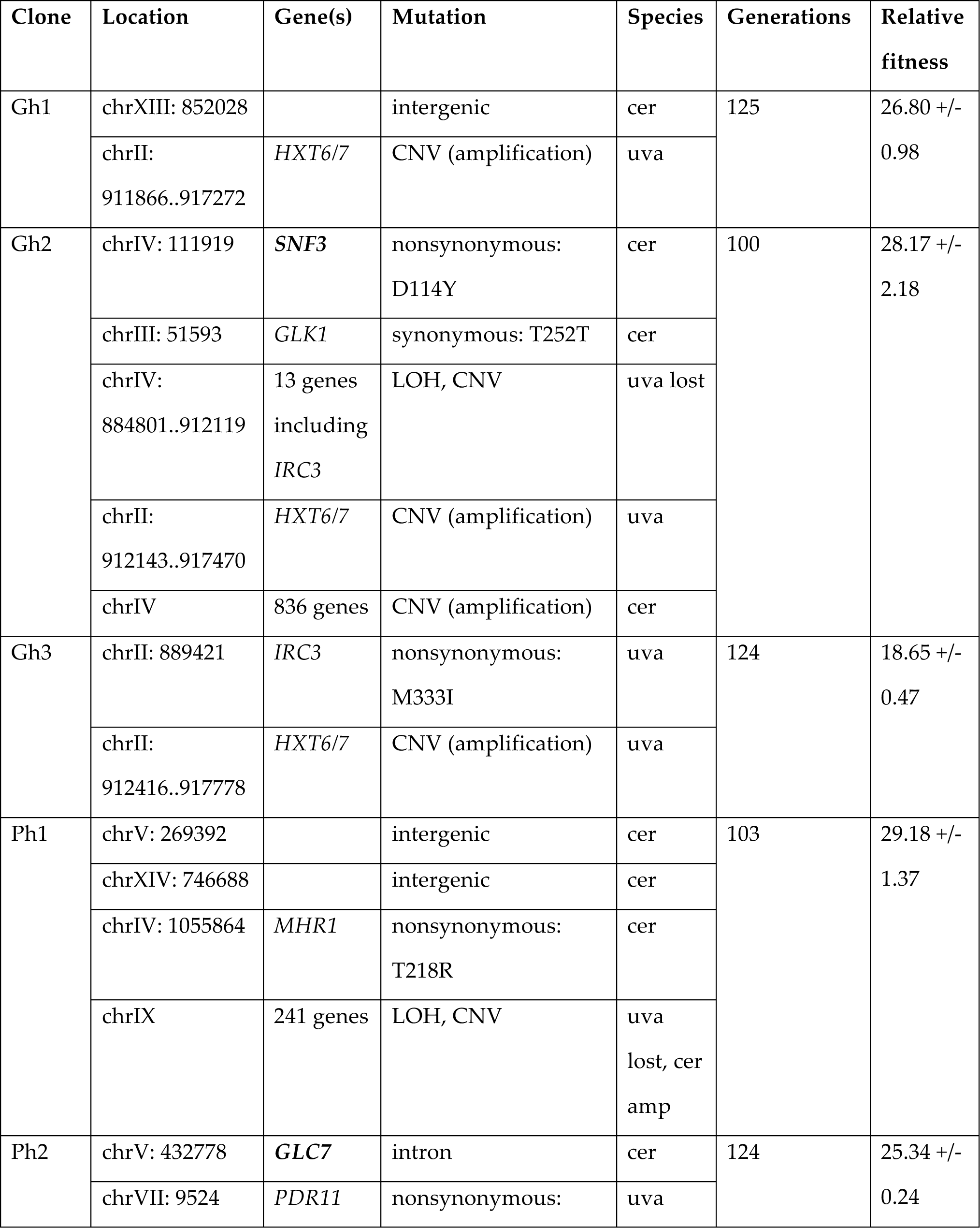
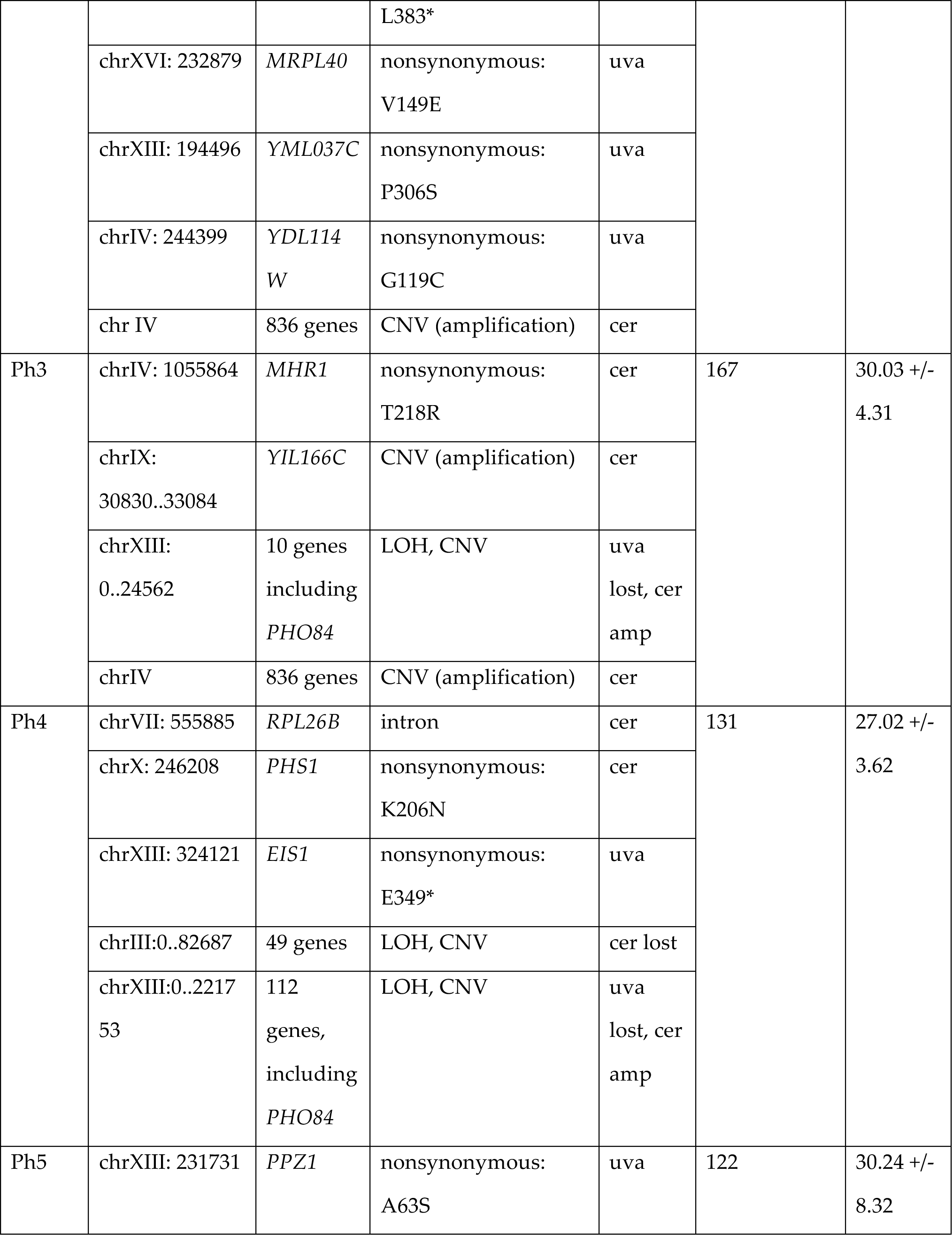
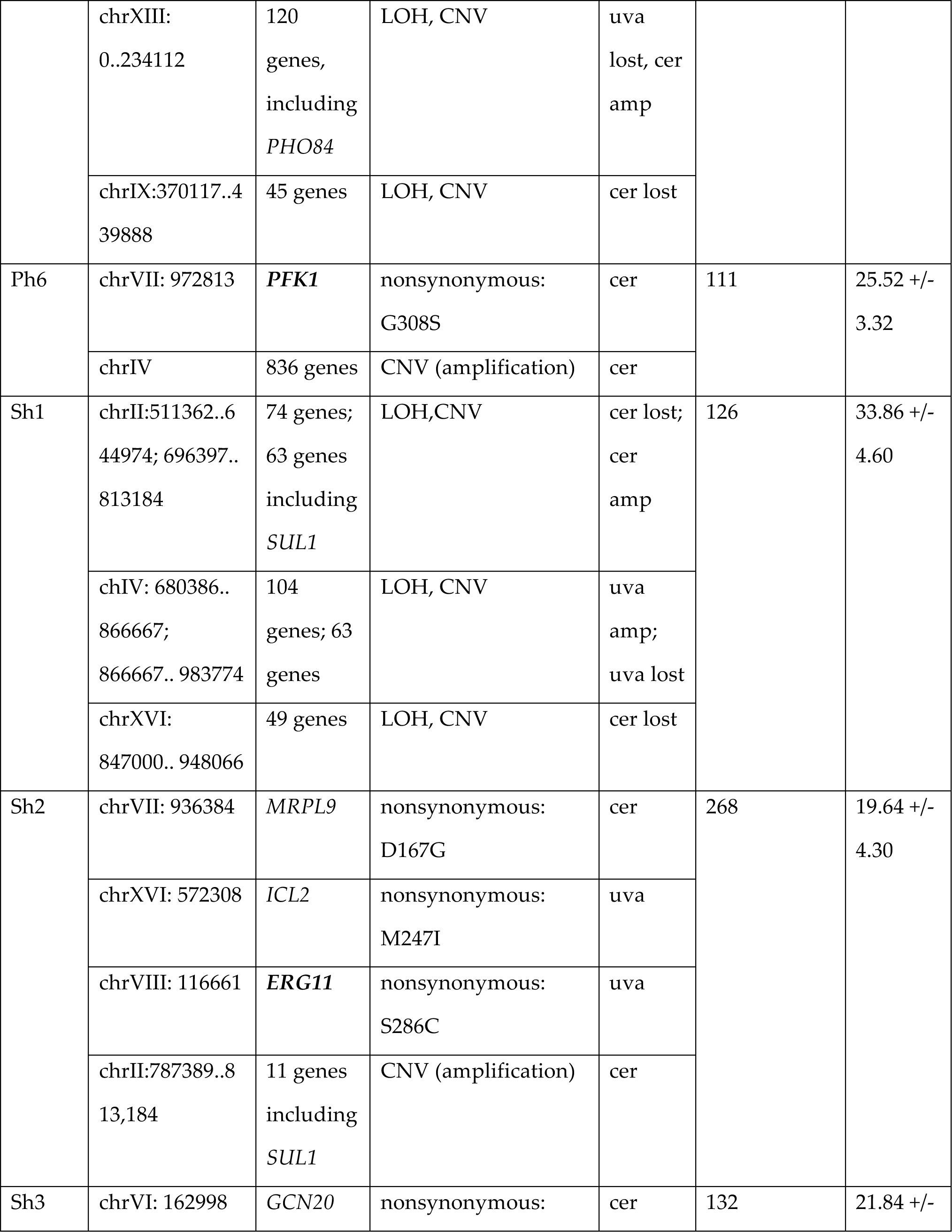
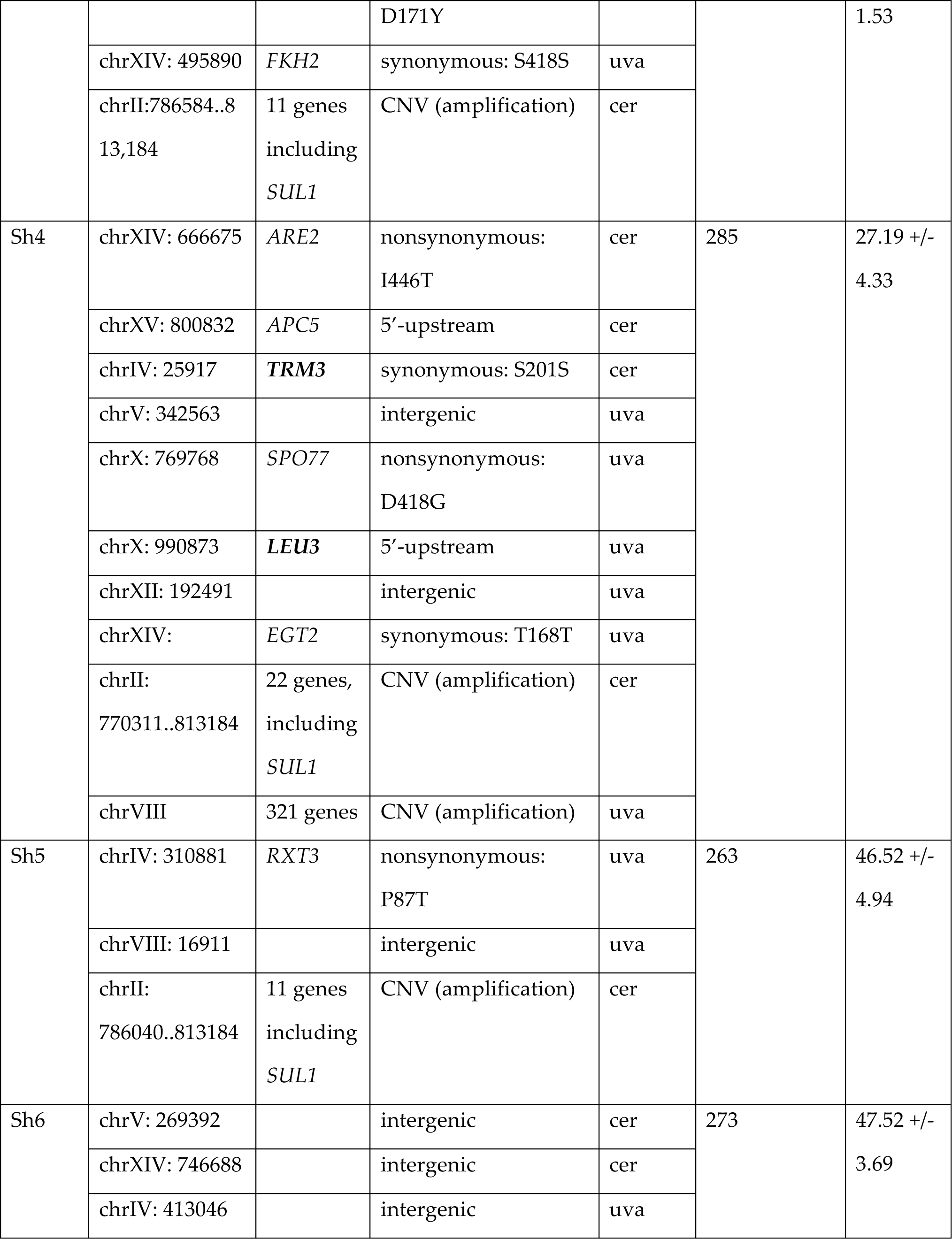
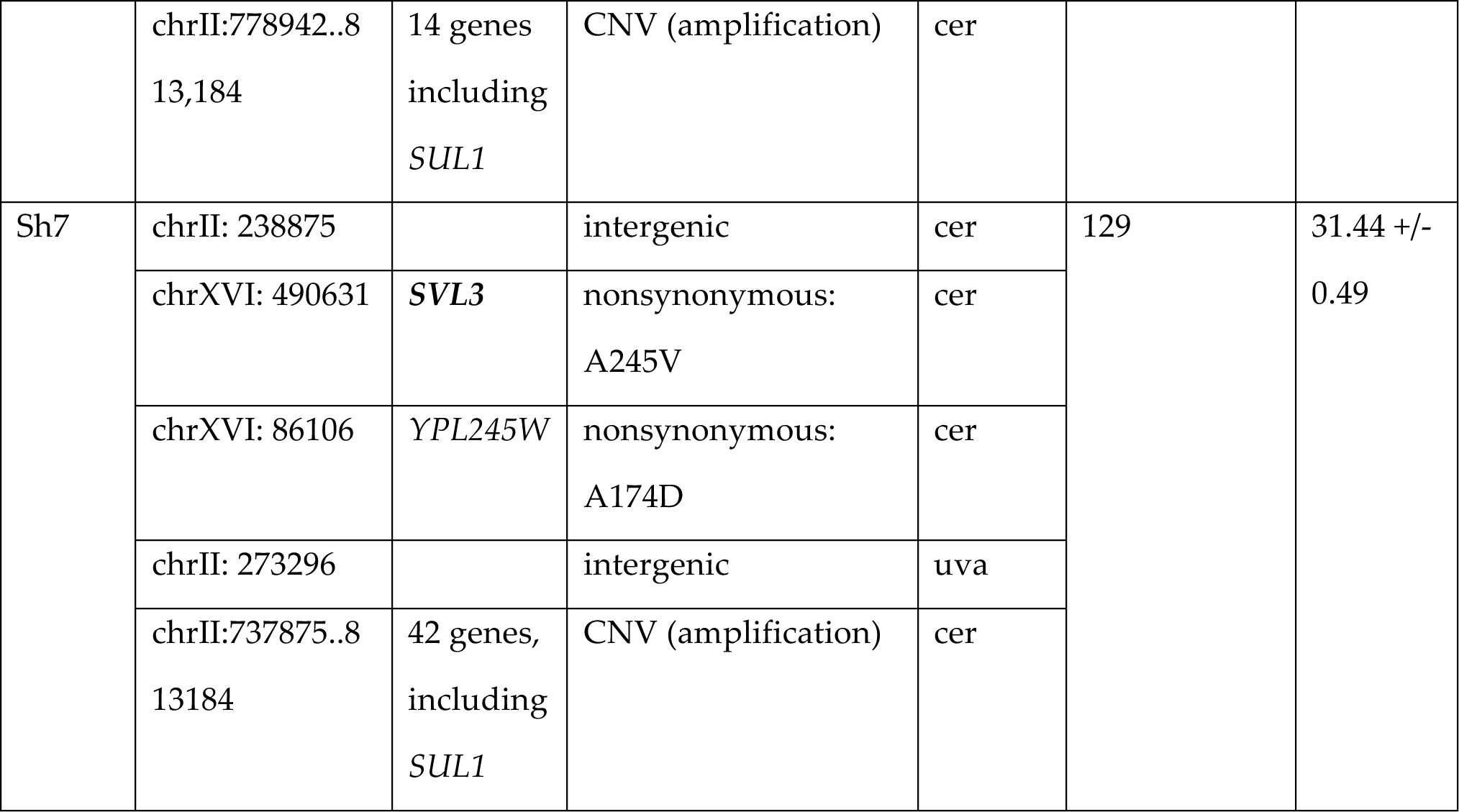
Mutations and fitness of evolved hybrid clones Clone Location. Point mutations, copy number variants (CNV), and loss of heterozygosity events (LOH) are recorded for each evolved hybrid clone. Clones are identified by nutrient (G: glucose-limitation, P: phosphate-limitation, and S: sulfate-limitation), an “h” denotes hybrid, and the number indicates its derivation from independent populations. Genes in bold have been found to have point mutations in prior experiments. Note that mutations in the *S. uvarum* genome use *S. uvarum* chromosomes and coordinates. All breakpoints were called by visual inspection of sequencing reads and are thus approximate.

**Table 2:**
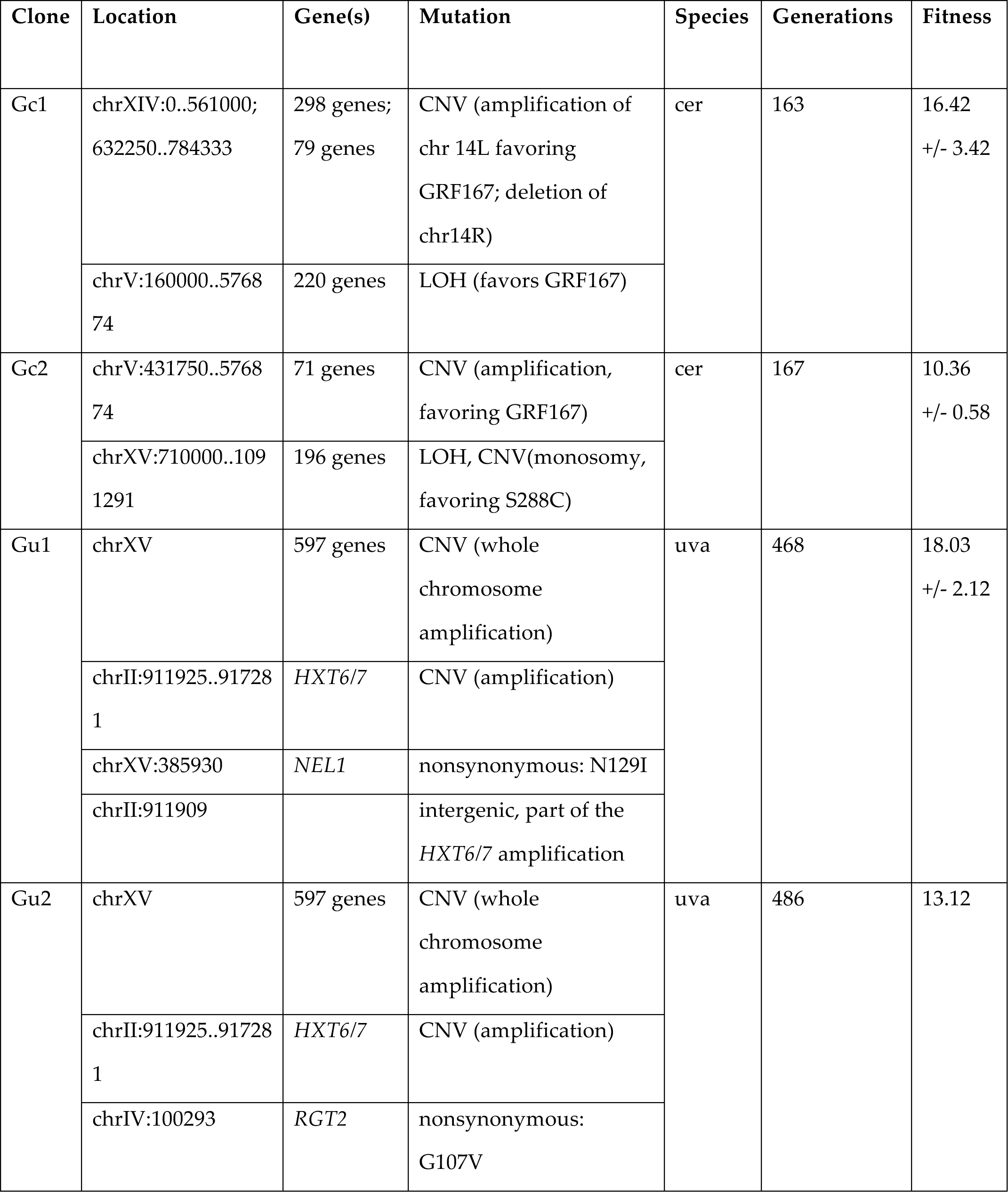
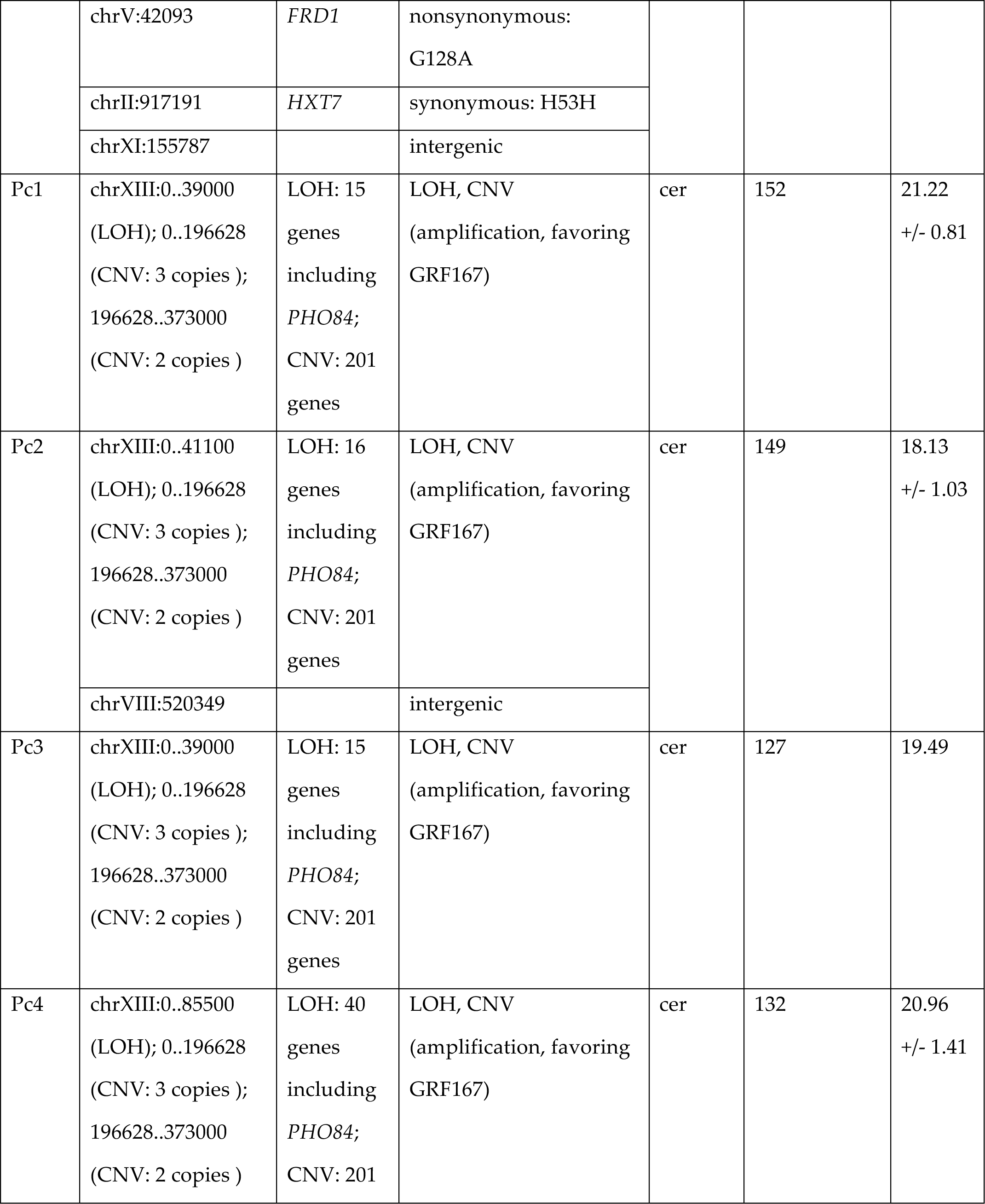
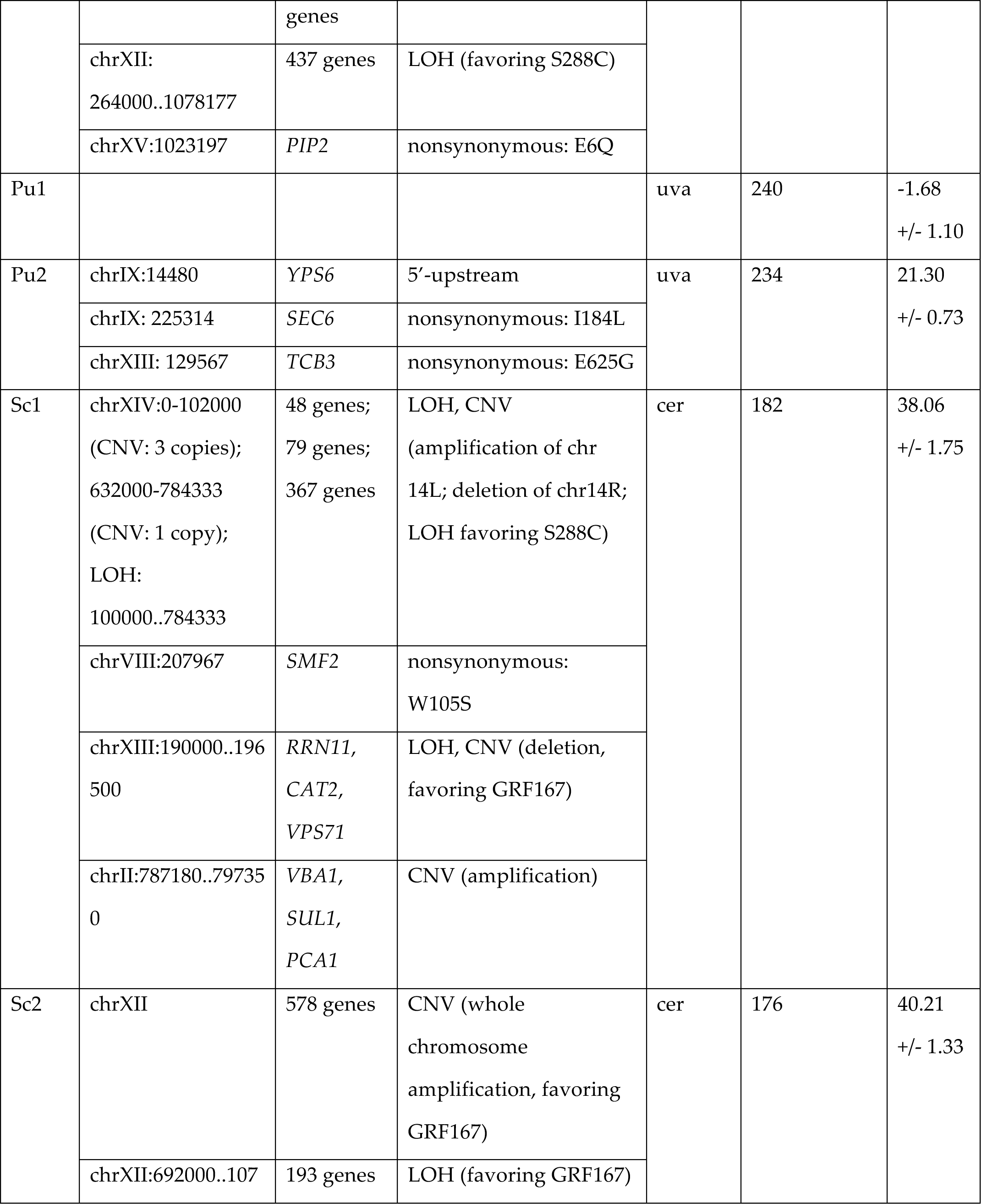
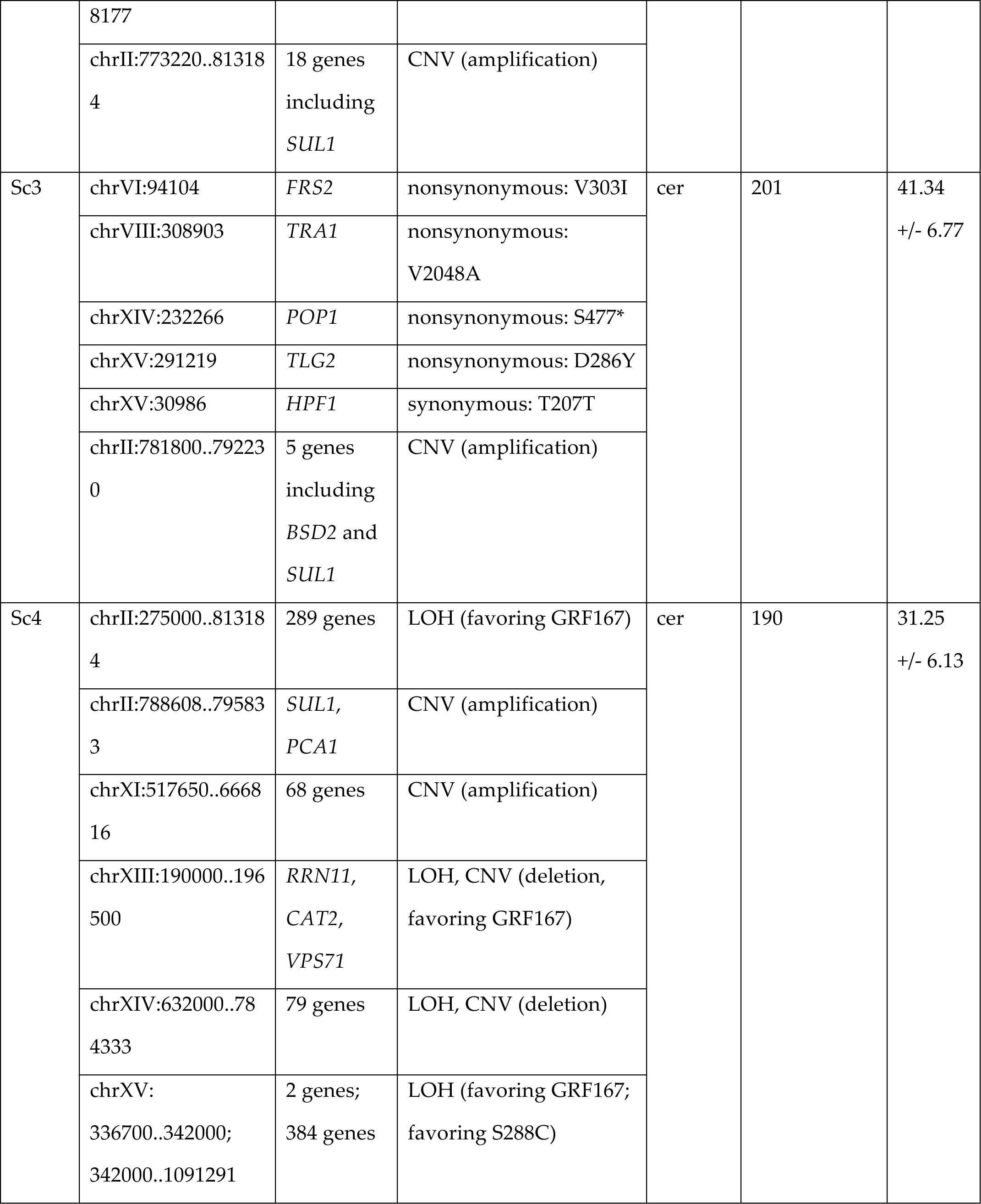
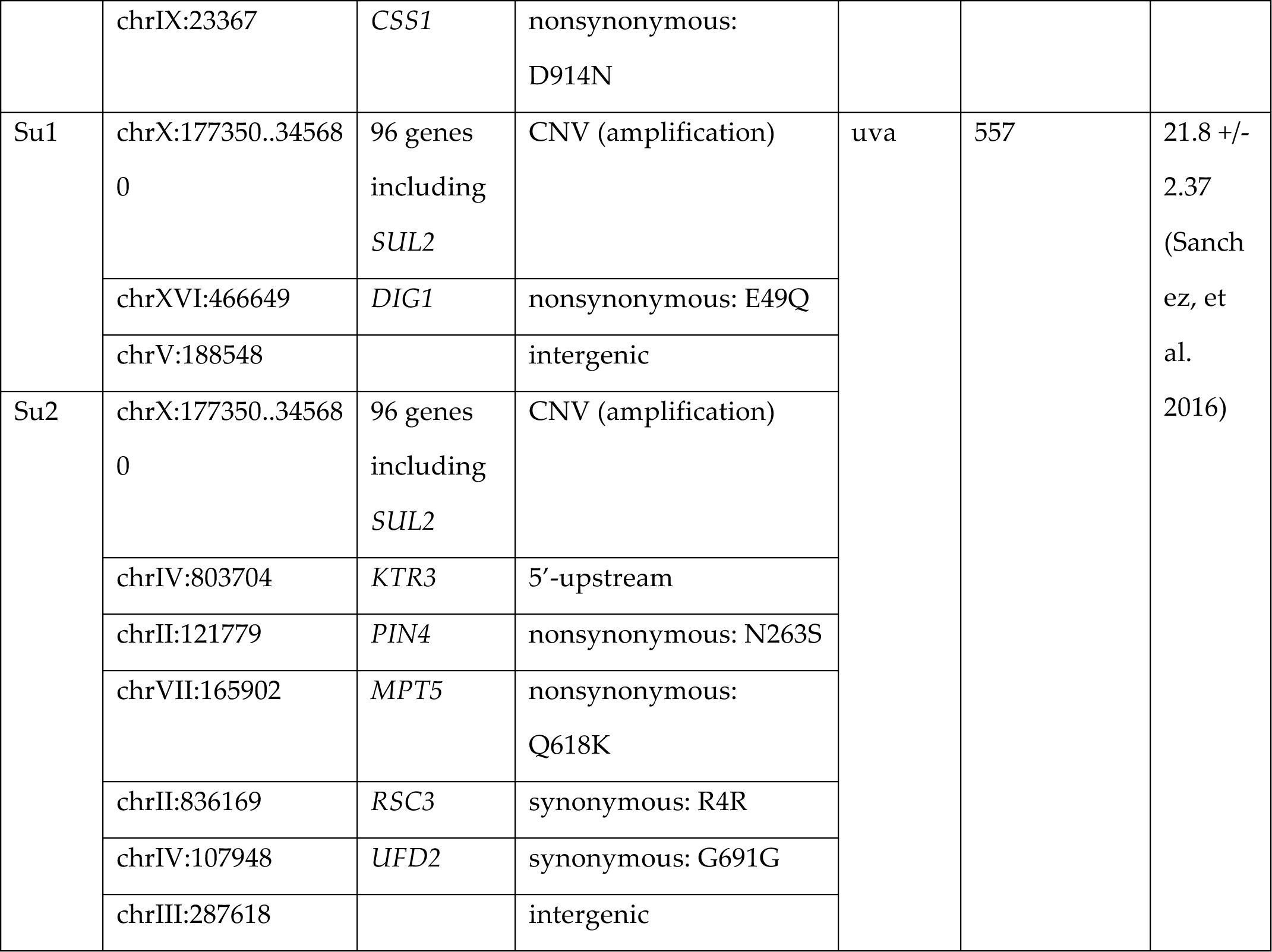
Mutations and fitness of evolved parental clones Clone Location. Point mutations, copy number variants (CNV), and loss of heterozygosity events (LOH) are recorded for each evolved parental clone. Clones are identified by nutrient (G: glucose-limitation, P: phosphate-limitation, and S: sulfate-limitation), by species (“c” denotes *S. cerevisiae*, “u” denotes *S. uvarum*), and the number indicates its derivation from independent populations. Note that mutations in the *S. uvarum* genome use *S. uvarum* chromosomes and coordinates. All breakpoints were called by visual inspection of sequencing reads and are thus approximate.

### Mutations in nuclear encoded mitochondrial genes may be more prevalent in interspecific hybrids

To identify mutations in the evolved hybrids, we generated whole genome sequencing for sixteen clones from the endpoints of the evolution experiments (**Table 1**). We thus captured data from a range of nutrient limitations (6-phosphate; 3-glucose; 7-sulfate) and generations (100-285, median 154 generations). Each clone had an average of 2.4 point mutations, a number of which have been previously identified in prior *S. cerevisiae* evolution experiments. For example, a nonsynonymous mutation in the *S. cerevisiae* allele of the glucose sensing gene *SNF3* has been identified in glucose limited experiments in *S. cerevisiae* (Kvitek and Sherlock 2013; Selmecki, et al. 2015). To our knowledge, 20/27 coding point mutations are unique to these experiments (Payen, et al. 2015).

In evolved parentals, we again sequenced one clone from the endpoint of each population. In total, we sequenced 16 clones, 6 from each of the three nutrients (two *S. uvarum* diploids, and four *S. cerevisiae* diploids), except in glucose limitation in which only two *S. cerevisiae* populations were sampled. The generations ranged from 234-557 (median 477) in *S. uvarum* with an average of 2.83 mutations per clone, and from 127-190 (median 166.5) in *S. cerevisiae* with an average of 0.9 point mutations per clone (**Table 2**). This discrepancy in point mutations between *S. cerevisiae* and *S. uvarum* may be explained by differences in generation time, or perhaps other mutational events are more prevalent in *S. cerevisiae*.

With the limited number of samples we have from hybrid and parental clones, it is difficult to draw any conclusions regarding unique point mutations in hybrids. However, one class of mutations that may be of particular interest in hybrids are genomic mutations which may interact with the mitochondria, as previous work has shown that nuclear-mitochondria interactions can underlie hybrid incompatibility (Chou and Leu 2010; Lee, et al. 2008; Meiklejohn, et al. 2013). Other studies have found that only the *S. cerevisiae* mitochondria are retained in *S. cerevisiae* x *S. uvarum* hybrids (Antunovics, et al. 2005), and we recapitulate these findings, potentially setting the stage for conflicting interactions between the *S. uvarum* nuclear genome and the foreign mitochondria. We observe several mitochondria related mutations in hybrids in both *S. cerevisiae* and *S. uvarum* alleles. For example, one point mutation, a non-synonymous mutation in the *S. cerevisiae* allele of the mitochondrial ribosomal protein gene *MHR1*, was seen in two separate clones independently evolved in phosphate limitation. This gene may be of particular interest as it was discovered in a previous screen as being haploproficient (increased fitness of 19%) in hybrids in which the *S. cerevisiae* allele is missing and the *S. uvarum* allele is retained (Lancaster S, Dunham MJ, unpublished data), suggesting that this mutation may alter or disable the *S. cerevisiae* protein in some way. Another example involves the gene *IRC3*, a helicase responsible for maintenance of the mitochondrial genome, which has a nonsynonymous mutation in the *S. uvarum* allele in clone Gh3 and is deleted in clone Gh2, potentially suggesting that the *uvarum* allele is deleterious in the hybrid background. While our sample size is small, 4/27 point mutations in hybrids are related to mitochondria function compared to 0/26 in parentals, and may represent interesting targets for further exploration.

### Copy number variants frequently involve the amplification of nutrient transporters

Yeast in both natural and artificial environments are known to frequently experience changes in copy number, ranging from single genes to whole chromosomes (Dunham, et al. 2002; Dunn, et al. 2012; Gresham, et al. 2008; Kvitek and Sherlock 2013; Payen, et al. 2014; Selmecki, et al. 2015; Sunshine, et al. 2015; Zhu, et al. 2016). This holds true in our evolution experiments: we observe copy number changes across all genetic backgrounds (**Figure 1**, **Supplemental Figures 1-3**). Clones were compared to array Comparative Genomic Hybridization of populations to confirm that clones are representative of populations (see Materials and Methods). The evolved hybrid clones displayed an average of 1.5 copy number variants (CNVs) per clone (**Figure 1**, **Table 1**, **Supplemental Figure 3**), as defined by the number of segmental or whole chromosome amplifications/deletions (though it is likely that some of these CNVs were created in the same mutational event). The evolved *S. cerevisiae* clones had an average of 1.5 CNV per clone and the evolved *S. uvarum* had an average of 1 CNV per clone (**Table 2**, **Supplemental Figures 1-2**). It therefore does not appear that our interspecific hybrids are more prone to genomic instability, as has previously been suggested in other systems (Chester, et al. 2015; Lloyd, et al. 2014; Mason and Batley 2015; Xiong, et al. 2011). The most common event across nutrient limitations in the interspecific hybrids was an amplification of the *S. cerevisiae* copy of chromosome IV, which occurred in four independent hybrid clones (3 in phosphate limitation, 1 in glucose limitation; **Supplemental Figure 3**). Several other characteristic rearrangements occurred in the evolved *S. cerevisiae* clones, including the amplification of the left arm of chromosome 14 accompanied by segmental monosomy of the right arm of chromosome 14, an event seen previously in other evolved populations (Dunham, et al. 2002; Gresham, et al. 2008; Sunshine, et al. 2015). All copy number events in *S. cerevisiae* had breakpoints at repetitive elements known as Ty elements, except those located on chrII, which are known to be mediated by another mechanism (Brewer, et al. 2015). In contrast, copy number variants in the hybrid were rarely facilitated by repetitive elements, perhaps in part because *S. uvarum* has no full length Ty elements.

**Figure 1:**
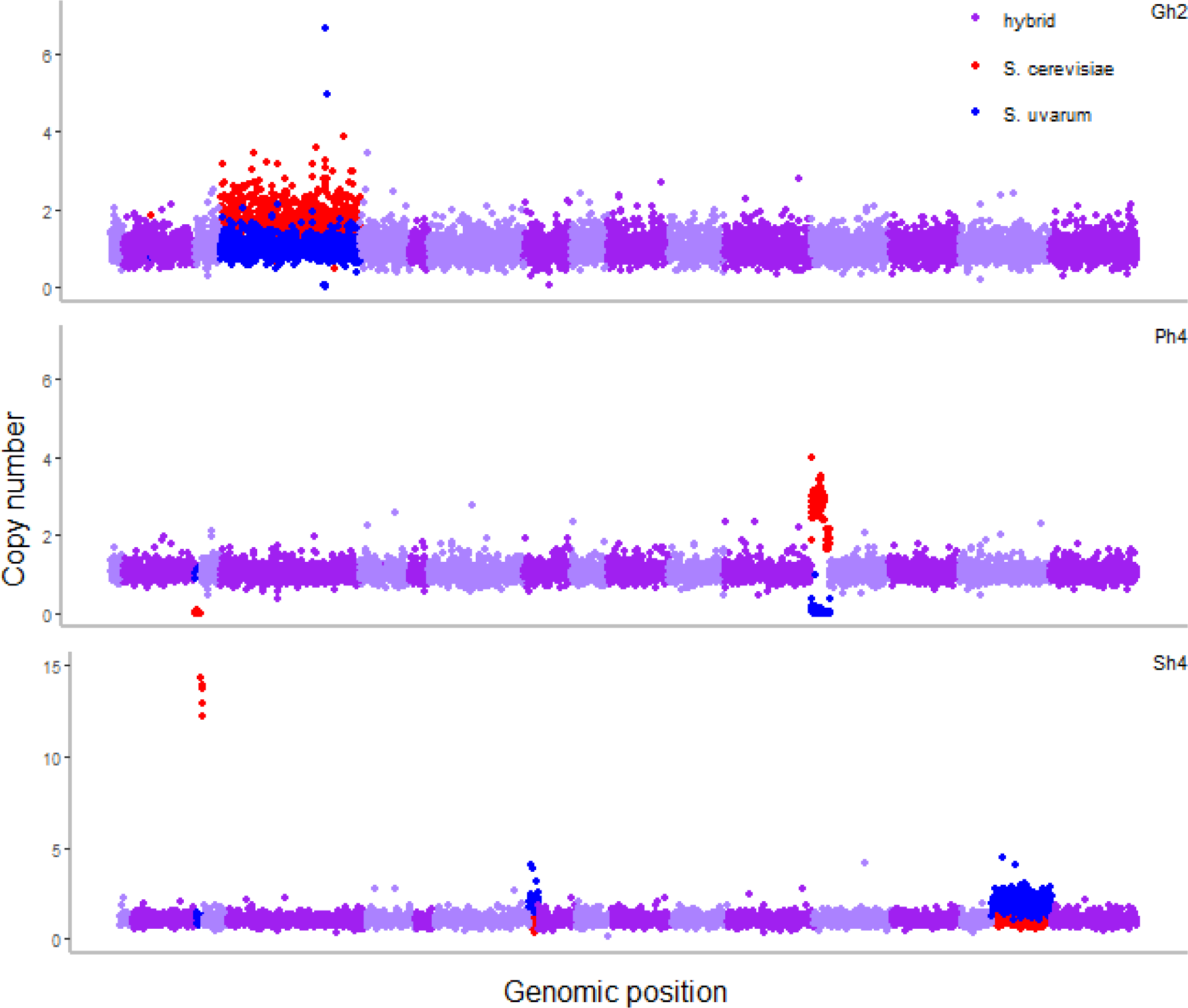
Evolved hybrids exhibit changes in copy number and loss of heterozygosity. Copy number variants are displayed for evolved hybrid clones from three nutrient limited conditions: Gh2, glucose; Ph4, phosphate; and Sh4, sulfate. Hybrid copy number, determined by normalized sequencing read depth per ORF, is plotted across the genome according to *S. cerevisiae* ORF coordinates to account for three reciprocal translocations between *S. cerevisiae* and *S. uvarum*. Chromosomes are plotted in alternating light and dark purple, red indicates a *S. cerevisiae* copy number variant and blue indicates a *S. uvarum* copy number variant. Gh2 has a whole chromosome amplification of *S. cerevisiae* chrIV, a small segmental deletion of *S. uvarum* chrIV (non-copy neutral loss of heterozygosity), and an amplification of *S. uvarum HXT6/7*. Ph4 has a small segmental deletion of *S. cerevisiae* chrIII (non-copy neutral loss of heterozygosity) and an amplification of *S. cerevisiae* chrXIII with corresponding deletion of *S. uvarum* chrXIII (copy neutral loss of heterozygosity). Sh4 has an amplification of *S. cerevisiae SUL1* and a whole chromosome amplification of *S. uvarum* chrVIII, (note, there is a reciprocal translocation between chrVIII and chrXV). Note that Sh4 is plotted on a different scale. For specific coordinates of copy number variants, see Table 1.

Frequently in nutrient limited evolution experiments, copy number variants involve amplification of the nutrient specific transporter, and indeed, we also observed amplification of these transporters in many of the clones. In sulfate limitation, the *S. cerevisiae* allele of the high affinity sulfate transporter *SUL1* is amplified in 7/7 hybrid clones and 4/4 *S. cerevisiae* clones (**Figure 1**, **Tables 1**-**2**, **Supplemental Figures 1**, **3**). Interestingly, *SUL2* is the preferred sulfate transporter in *S. uvarum* (Sanchez, et al. 2016) and was not observed to be amplified in the evolved hybrids (**Supplemental Figure 2**, **Table 2**). In glucose limitation, previous *S. cerevisiae* evolution experiments found consistent amplification of the high affinity glucose transporter genes *HXT6/7* (Brown, et al. 1998; Dunham, et al. 2002; Gresham, et al. 2008; Kao and Sherlock 2008; Kvitek and Sherlock 2011). In our experiments, the *S. uvarum* alleles of the *HXT6/7* transporters are amplified in 3/3 hybrid clones and both *S. uvarum* clones, but are not amplified in evolved *S. cerevisiae* clones, suggesting that the *S. uvarum HXT6/7* alleles confer a greater fitness advantage compared to *S. cerevisiae* (**Figure 1**, **Tables 1**-**2**, **Supplemental Figures 1-3)**. Finally, in phosphate limitation, the *S. cerevisiae* copy of the high affinity phosphate transporter *PHO84* is amplified, while the *S. uvarum* allele is lost in 3/6 hybrid clones in an event known as loss of heterozygosity (**Figures 1**-**2**, **Table 1**, **Supplemental Figure 3**). Intriguingly, the evolved *S. cerevisiae* also display loss of heterozygosity and accompanied amplification favoring the allele derived from strain GRF167 over the S288C allele in 4/4 clones (**Figure 2**, **Table 2**). All hybrid clones carry the “preferred” GRF167 *S. cerevisiae* allele, as this was the allele used to create the *de novo* hybrid.

**Figure 2:**
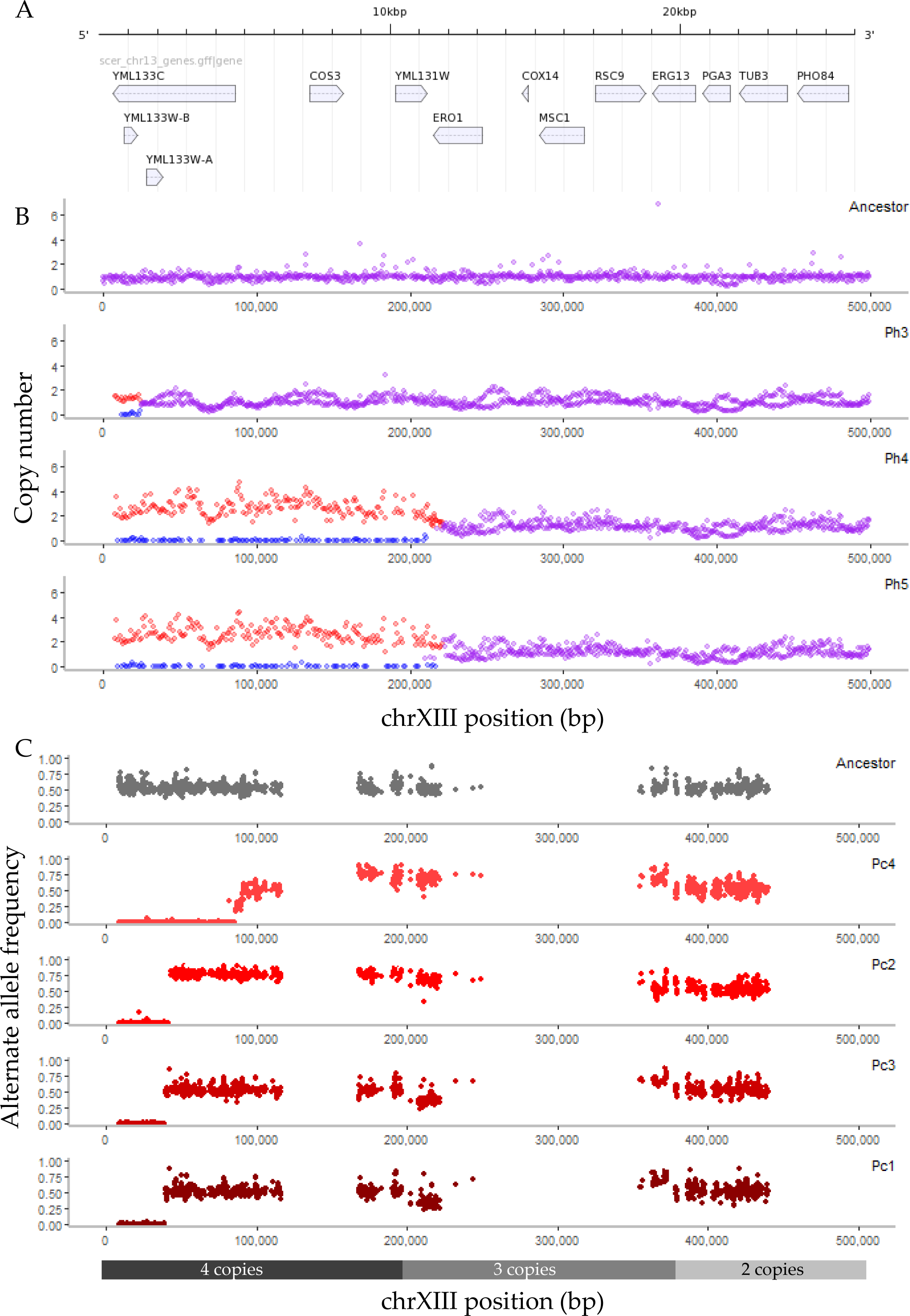
Repeated loss of heterozygosity at the *PHO84* locus in intra- and interspecific hybrids. A. The 25kb region extending from the left telomere of chromosome XIII to the high affinity phosphate transporter gene *PHO84*. B. Copy number is plotted across the whole chromosome XIII in the hybrid ancestor and three evolved hybrid clones in phosphate limitation (clone indicated in upper right corner). Red shows the *S. cerevisiae* allele, blue shows the *S. uvarum* allele, and purple shows where both species exhibit the same copy number. Note: 8kb of telomere sequence is removed due to repetitive sequence. C. Alternate allele frequency is plotted for a portion of chromosome XIII in the ancestor and four evolved *S. cerevisiae* clones in phosphate limitation (clone indicated in upper right corner). All evolved *S. cerevisiae* clones exhibit a loss of heterozygosity at the telomeric portion of chromosome XIII (loss of S288C, amplification of GRF167), as illustrated by an allele frequency of zero compared to the ancestor. *S. cerevisiae* copy number for the four evolved clones is shown below; the ancestor is diploid across the chromosome (also see **Table 2**, **Supp. Fig. 1**).

### Loss of heterozygosity is a common event in heterozygous evolving populations

Selection on heterozygosity, as a loss of heterozygosity event could represent, is an underappreciated source of adaptation in microbial experimental evolution, as typical experiments evolve a haploid or homozygous diploid strain asexually and as a result, have little variation to select upon. Loss of heterozygosity is observed in natural and industrial hybrids (Albertin and Marullo 2012; Louis, et al. 2012a; Pryszcz, et al. 2014a; Wolfe 2015), but here we document its occurrence in both intra- and interspecific hybrids in the laboratory as a result of short term evolution (also see (Burke, et al. 2014; Dunn, et al. 2013)). Loss of heterozygosity is observed across all nutrient conditions, with twelve independent loss of heterozygosity events detected in *S. cerevisiae*, and nine independent events documented in the hybrids (**Figures 1**-**2**, **Tables 1**-**2**, **Supplemental Figures 1, 3**). It thus appears that this type of mutational event is both common, and can occur over short evolutionary timescales.

The loss of heterozygosity event can result in copy-neutral (where one allele is lost and the other allele is amplified) or non-neutral chromosomal segments (where one allele is lost, rendering the strain hemizygous at that locus), and can favor the retention of either allele. In *S. cerevisiae*, there is a bias in resolution where loss of heterozygosity events favor retaining the GRF167 allele over the S288C allele (**Table 2**, **Supplemental Figure 4**). One unique case in clone Sc4 has a small ∼5 kb loss of heterozygosity event on chrXV favoring GRF167, which switches to favoring S288C for the rest of the chromosome. The retention of *S. cerevisiae* is slightly more common in the hybrids (5/9 events, **Table 1**, **Supplemental Figure 3**), though not as drastic as the observed genome resolution in the hybrid *S. pastorianus*, where loss of heterozygosity favors *S. cerevisiae* over *S. eubayanus* (Nakao, et al. 2009). The size of the event ranges from approximately 25 kb to the whole chromosome level in the hybrids, and from 5kb to 540kb in *S. cerevisiae*. Where loss of heterozygosity is accompanied by an amplification event, the loss of heterozygosity event always occurs first; unlike many CNV events, almost all loss of heterozygosity events do not appear to be mediated by existing repetitive sequence such as a transposable element in the hybrid or *S. cerevisiae*, and are most likely a product of break induced replication or mitotic gene conversion (Hoang, et al. 2010). The exceptions are in hybrid clones Ph4, Ph5, and Sh1, where there is a non-copy neutral loss of *S. cerevisiae* mediated by a Ty element or Long Terminal Repeat (LTR), and *S. cerevisiae* clones Sc1 and Sc4, where there is a 6.5 kb deletion of the S288C allele flanked by two Ty elements.

Loss of heterozygosity events in hybrids could signify several ongoing processes in hybrid genome evolution: loss of heterozygosity regions may represent (1) loci with incompatibilities; (2) selection on existing variation; or (3) genetic drift eroding genomic segments. While our sample size is modest, failing to see repeated loss of heterozygosity events across nutrient conditions disfavors the hypothesis that loss of heterozygosity is resolving some sort of hybrid incompatibility. Furthermore, loss of heterozygosity events observed in evolved *S. cerevisiae* suggest that this mutation type is not unique to interspecific hybrids. Instead, repeated events within a particular condition, such as the repeated loss of heterozygosity at *PHO84* in phosphate limitation or the 6.5kb segment on chrXIII in sulfate limitation, suggest that these events are beneficial, and are indeed selection on one allele over the other.

### Loss of heterozygosity is driven by selection on one allele

To test the hypothesis that loss of heterozygosity events provide a selective advantage, we used allele replacement, in which the allele of one species/strain is replaced with the allele of the other species/strain in an otherwise isogenic background. We tested this hypothesis using the most commonly seen loss of heterozygosity event, loss of heterozygosity at *PHO84*. While the region extends from 25-234kb in length in the hybrids and 40-85 kb in *S. cerevisiae*, *PHO84* was a prime candidate driving this event. *PHO84* is one of only 10 genes encompassed in the region extending from the telomere to the breakpoint of the shortest loss of heterozygosity event, and is included in every other loss of heterozygosity event on chromosome XIII (Figure 1). It is responsible for sensing low phosphate, and previous work identified a point mutation in *PHO84* (an Alanine to Valine substitution at the 5’ end of the gene), which increased fitness by 23% in phosphate limited conditions (Sunshine, et al. 2015). Finally, prior work with other nutrient transporters has shown amplification of nutrient transporters to be a key event in adapting to nutrient limited conditions.

We thus selected a region of approximately 2.5kb encompassing the *PHO84* ORF, its promoter, and 3’UTR (Cherry, et al. 2012; Nagalakshmi, et al. 2008; Yassour, et al. 2009). We created allele replacement strains using the two alleles of *S. cerevisiae* in a *S. cerevisiae* diploid background; the two alleles are 99.1% identical in this region and each strain is identical to the ancestral strain used in our evolution experiments except at the *PHO84* locus. The *S. cerevisiae* ancestor carries one copy of GRF167 (“preferred”) and one copy of S288C (“un-preferred”), so named due to which allele was retained and amplified in the evolved clones. To measure any resultant changes in fitness, we competed each strain individually against a fluorescent ancestral strain and measured which strain overtook the culture. Two copies of the un-preferred allele decreased fitness by -5.31 (+/-1.86), while two copies of the preferred allele increased fitness by 9.93 (+/-0.27). This displays an overall difference in fitness of 15.24 between the un-preferred and preferred alleles. By comparing the fitness of these allele replacement strains to the evolved clones (**Table 2**), the allele replacement does not fully recapitulate the fitness gain observed in the evolved clone. One explanation is that the additional mutations present in the evolved strains also contribute to their total fitness. Another explanation could be the increased copy number of the *PHO84* region that we see in these evolved clones. To further explore this fitness difference, we cloned the GRF167 allele onto a low copy number plasmid and transformed the allele replacement strain carrying two preferred *S. cerevisiae* alleles to simulate increased copy number of *PHO84*, and saw a further fitness increase of 1.76. This supports the conclusion that relative fitness gains in the evolved clone are largely due to the loss of the S288C allele, and selection and amplification of the GRF167 allele, with little additional benefit from further amplification. It could also be the case that co-amplification of other genes in the segment is required to attain the full benefit, as previously observed by the contribution of *BSD2* to the *SUL1* amplicon (Sunshine, et al. 2015; Payen, et al. 2015).

Unfortunately, we were unable to obtain a successful strain carrying the preferred *S. cerevisiae* allele in an *S. uvarum* background, so we were unable to test the fitness effect of carrying two preferred *S. cerevisiae* alleles in the hybrid background (which typically carries one preferred *S. cerevisiae* allele and one *S. uvarum* allele). However, we were able to generate a *S. uvarum* strain carrying the unpreferred allele and use this to create a hybrid. Carrying one preferred allele and one un-preferred *S. cerevisiae* allele has an increased fitness of 4.35 compared to two *S. uvarum* alleles in a hybrid background. Furthermore, using the GRF167 *PHO84* plasmid, we found that the hybrid has an increased fitness of 16.47 (+/- 1.13) when an extra preferred allele is added. Together, these results support the conclusion that the *S. cerevisiae* GRF167 allele is preferred over the S288C one, and that *S. cerevisiae* alleles are preferred over the *S. uvarum* allele in the hybrid, and hence, that the loss of heterozygosity events seen in both intra- and interspecific hybrids are the product of selection.

## DISCUSSION

In summary, we sought to understand forces underlying genome stabilization and evolution in interspecific and intraspecific hybrids as they adapt to novel environments. We evolved and sequenced clones from 16 hybrid populations and 16 parental populations to reveal a variety of mutational events conferring adaptation to three nutrient limited conditions. Of particular note, we find loss of heterozygosity in both evolved intraspecific and interspecific hybrid clones in all nutrient environments, potentially signifying areas where selection has acted on preexisting variation present in the ancestral clone. We used an allele replacement strategy to test this hypothesis for a commonly repeated loss of heterozygosity event and show that selection is indeed driving the homogenization of the genome at this locus. Though other studies in natural, industrial, and lab-evolved isolates have observed loss of heterozygosity, we present the first empirical test of the causal evolutionary forces influencing these events. This work is particularly informative for understanding past hybridization events and subsequent genome resolution in hybrids in natural and artificial systems.

### The predictability of evolution

We now have many examples of predictable evolution in natural systems (Conte, et al. 2012; Elmer and Meyer 2011; Jones, et al. 2012; Losos, et al. 1998; Martin and Orgogozo 2013; Rundle, et al. 2000; Wessinger and Rausher 2014), and in laboratory experimental evolution, in which there often appears to be a limited number of high fitness pathways that strains follow when adapting to a particular condition (Burke, et al. 2010; Ferea, et al. 1999; Gresham, et al. 2008; Kawecki, et al. 2012; Kvitek and Sherlock 2013; Lang and Desai 2014; Salverda, et al. 2011; Woods, et al. 2006). For example, it is well established that amplifications of nutrient transporters are drivers of adaptation in evolution in nutrient limited conditions. Previous work in our group has particularly focused on the amplification of the high affinity sulfate transporter gene *SUL1* in sulfate limited conditions, which occurs in almost every sulfate limited evolution experiment and confers a fitness advantage of as much as 40% compared to its unevolved ancestor. The amplification of phosphate transporters has been markedly less common, and thus drivers of adaptation in this condition have been less clear. Gresham et al. (2008) identified a whole chromosome amplification of chrXIII in one population. In a follow up study, Sunshine et al. (2015), found whole or partial amplification of chrXIII in 3/8 populations. A genome wide screen for segmental amplifications found a slight increase in fitness for a small telomeric segment of chromosome XIII, and a point mutation in *PHO84* was observed to increase fitness by 23%. However, screens by Payen *et al.* (2015) showed that although *PHO84* is recurrently mutated in various experiments, it showed no benefit when amplified or deleted in phosphate limited conditions. Finally, additional evolution experiments recapitulated the point mutation seen in Sunshine et al. in 24/32 populations, and saw amplification of *PHO84* in 8/32 populations (Miller A, Dunham MJ, unpublished data). It is important to note that all of these experiments used a strain background derived from S288C or CEN.PK, both of which carry the same *PHO84* allele.

In our work, we observed amplification of the *S. cerevisiae* GRF167 allele of *PHO84* in 4/4 *S. cerevisiae* clones from 4 populations and 3/6 hybrid clones from 6 populations. This amplification was always preceded by the loss of the S288C allele in *S. cerevisiae* clones or occurred in conjunction with the loss of the *S. uvarum* allele in hybrids. Furthermore, there is a 15% fitness difference between carrying two copies of the S288C allele of *PHO84* compared to carrying two copies of the GRF167 allele of *PHO84*. It thus appears that amplification of *PHO84* has been less predictable as the S288C allele does not confer a fitness advantage unless mutated. We note that the preferred GRF167 allele of *PHO84* does not carry this particular polymorphism. Together, these results imply that strain background can constrain adaptive pathways.

The infusion of variation created by hybridization provides new templates for selection to act upon, which can be more important than either point mutations or copy number variants alone. Our work shows that outcrossing need not be common to have long-lasting effects on adaptation. This implication is particularly relevant in yeast where outcrossing may occur quite rarely followed by thousands of asexual generations (Greig and Leu 2009; Liti 2015; Ruderfer, et al. 2006).

### Applications to other hybrids and cancer

The observation that loss of heterozygosity occurs in hybrid genomes is increasingly documented (Borneman, et al. 2014; Louis, et al. 2012b; Marcet-Houben and Gabaldon 2015; Pryszcz, et al. 2014b; Soltis, et al. 2014), although the reason(s) for this type of mutation has been unresolved. As most examples stem from allopolyploid events that occurred millions of years ago, understanding why loss of heterozygosity is important in hybrid genome evolution is difficult. Cancer cells are also known to experience loss of heterozygosity, sometimes involved in the inactivation of tumor suppressor genes, leaving only one copy of the gene that may be mutated or silenced (Lapunzina and Monk 2011; Thiagalingam, et al. 2001; Tuna, et al. 2009). Data support the conclusion that loss of heterozygosity events are selected for during tumor development, as many loss of heterozygosity events involve specific chromosomal segments (Thiagalingam, et al. 2001), although the underlying molecular and genetic reasons for selection is an open debate (Ryland, et al. 2015).

Here, we experimentally demonstrate that loss of heterozygosity can occur in homoploid hybrids, as well as intraspecific hybrids. We provide an example in which homogenization of the genome is non-random, but instead driven by selection on one allele. We furthermore discover examples where one species allele is preferred over the other without loss of heterozygosity, such as the repeated amplification of the *S. uvarum* high affinity glucose transporters *HXT6/7*. Amplification of one species allele with or without loss of heterozygosity may be due to hybrid incompatibility within a particular protein complex, or other epistatic interactions (Piatkowska, et al. 2013). Together, our results show that the heterozygosity supplied by hybridization is an important contributor to adaptive routes explored by populations as they adapt to novel conditions.

While we cannot generalize our results from the *PHO84* locus across the many other loss of heterozygosity events discovered in our hybrids and *S. cerevisiae*, in the future we can use similar methodology to explore whether positive selection always drives loss of heterozygosity or whether other explanations such as incompatibility resolution contribute as well. Future experiments might also utilize a high throughput method to explore segmental loss of heterozygosity in hybrids at a genome wide scale, similar to ongoing experiments at the gene level (Lancaster S, Dunham MJ, unpublished data). While our sample size is modest, this is a novel and necessary step in understanding forces underlying hybrid genome stabilization and highlighting an underappreciated mechanism of hybrid adaptation.

### Conclusions

The mutation events we observe in our experimentally evolved hybrids are in many ways quite representative of mutations observed in ancient hybrid genomes, suggesting that hybrid genome stabilization and adaptation can occur quite rapidly (within several hundred generations). Furthermore, our results illustrate that the infusion of variation introduced by hybridization at both the intra- and inter-species level can increase fitness by providing choices of alleles for selection to act upon, even when sexual reproduction is rare. This may be particularly important for leveraging existing variation for agricultural and industrial processes, and as climate change potentially increases natural hybridization (Hoffmann and Sgro 2011; Kelly, et al. 2010; Muhlfeld, et al. 2014).

## MATERIALS AND METHODS

### Strains

A list of strains used in this study is included in **Supplemental Table 1**. All interspecific hybrids were created by crossing a *ura3 LYS2* haploid parent to a *URA3 lys2* haploid parent of the other mating type, plating on media lacking both uracil and lysine, and selecting for prototrophs.

### Evolution Experiments

Continuous cultures were established using media and conditions previously described (Gresham, et al. 2008; Sanchez, et al. 2016). Detailed protocols and media recipes are available at http://dunham.gs.washington.edu/protocols.shtml. Samples were taken daily and measured for optical density at 600 nm and cell count; microscopy was performed to check for contamination; and archival glycerol stocks were made daily. An experiment was ended when contamination, growth in tubing, or clumping appeared (number of generations at the endpoint for each population shown in **Tables 1**, **2**). Samples from each endpoint population were colony-purified to yield two clones for further study.

### Array Comparative Genomic Hybridization (aCGH)

Populations from the endpoint of each evolution were analyzed for copy number changes using aCGH following the protocol used in Sanchez *et al.* (2016). Microarray data will be made available upon publication in the GEO database and the Princeton University Microarray Database.

### Sequencing

DNA was extracted from overnight cultures using the Hoffman-Winston protocol (Hoffman and Winston 1987), and cleaned using the Clean & Concentrator kit (Zymo Research). Nextera libraries were prepared following the Nextera library kit protocol and sequenced using paired end 150 base pairs on the illumina NextSeq 500 machine (sequencing coverage in **Supplemental Table 2**). The reference genomes used were: *S. cerevisiae* v3 (Engel, et al. 2014), *S. uvarum* (Scannell, et al. 2011), and a hybrid reference genome created by concatenating the two genomes. Sequence was aligned to the appropriate reference genome using bwa v0.6.2 (Li and Durbin 2009) and mutations were called using GATK (McKenna, et al. 2010) and samtools 0.1.19 (Li, et al. 2009). Mutations in evolved clones were filtered in comparison to the ancestor to obtain *de novo* mutations. All mutations were first visually inspected using Integrative Genomics Viewer (Robinson, et al. 2011). Subsequently, point mutations in the hybrids were confirmed with Sanger sequencing (**Supplemental Table 3**). Copy number variants were visualized using DNAcopy for *S. cerevisiae* and *S. uvarum* (Seshan and Olshen 2016). Loss of heterozygosity events were called based on sequencing coverage in the hybrids, and by identifying homozygous variant calls in *S. cerevisiae*. All breakpoints were called by visual inspection of sequencing reads and are thus approximate.

### Fitness assays

The pairwise competition experiments were performed in 20 mL chemostats (Miller and Dunham 2013). Each competitor strain was cultured individually until steady state was reached, and then was mixed 50:50 with a GFP-tagged ancestor. Each competition was conducted in two biological replicates for approximately 15 generations after mixing. Samples were collected and analyzed twice daily. The proportion of GFP+ cells in the population was detected using a BD Accuri C6 flow cytometer (BD Biosciences). The data were plotted with ln[(dark cells/GFP+ cells)] vs. generations. The relative fitness coefficient was determined from the slope of the linear region.

### Strain construction

Allele replacements for the *PHO84* locus were done following the protocol of the Caudy lab with further modifications described here. The native locus was replaced with *Kluyveromyces lactis URA3*. The *pho84Δ::URA3* strain was grown overnight in 5 mL of C-URA media, then inoculated in a flask of 100 mL YPD and grown to an OD of 0.6-0.8. Cells were washed then aliquoted. 275 µl of transformation mix (35 µl 1M Lithium Acetate, 240 µl of 50% 3500 PEG), 10 µl of Salmon sperm, and approximately 3 µg of PCR product were added to the cell pellet. It was incubated at 37°C (*S. uvarum*) or 42°C (*S. cerevisiae*) for 45 minutes, then plated to YPD. It was replica plated to 5FOA the following day and colonies were tested for the gain of the appropriate species allele. The GRF167 allele was cloned into the pIL37 plasmid using Gibson assembly (Gibson, et al. 2009). Correct assembly was verified by Sanger sequencing. All primers used can be found in **Supplemental Table 3**.

## ACKNOWLEDGEMENTS

We thank Noah Hanson and Erica Alcantara for technical assistance, and Monica Sanchez for helpful comments on this manuscript. Thanks to Yixian Zheng and Doug Koshland for contributing to the initial experimental design, creating yeast strains, and purchasing the oligonucleotides used for the microarrays. This work was supported by the National Science Foundation (grant number 1516330) and the National Institutes of Health (grant number R01 GM094306). MD is a Senior Fellow in the Genetic Networks program at the Canadian Institute for Advanced Research and a Rita Allen Foundation Scholar. CSH was supported in part by National Institutes of Health (grant number T32 HG00035). This work was also supported by the National Institutes of Health (grant number P50 GM071508) to the Lewis-Sigler Institute and from the Howard Hughes Medical Institute to Doug Koshland and Yixian Zheng.

## SUPPLEMENTARY FIGURES

**Fig S1: Whole genome copy number variation in *S. cerevisiae* evolved clones**

Sequencing coverage for each evolved clone was normalized using the ancestor, and copy number variants were inferred by changes from a copy number of two using DNAcopy for each nutrient condition **A.** glucose-limitation, **B.** phosphate-limitation, and **C.** sulfate-limitation. Chromosomes are plotted in alternating grey and red. The average copy number is plotted as a black line.

**Fig S2: Whole genome copy number variation in *S. uvarum* evolved clones**

Sequencing coverage for each evolved clone was normalized using the ancestor, and copy number variants were inferred by changes from a copy number of two using DNAcopy for each nutrient condition **A.** glucose-limitation, **B.** phosphate-limitation, and **C.** sulfate-limitation. Chromosomes are plotted in alternating grey and blue. The average copy number is plotted as a black line. Note that plots use *S. uvarum* chromosomes and coordinates.

**Fig S3: Whole genome copy number variation in hybrid evolved clones**

Copy number as determined by normalized sequencing read depth per ORF is plotted across the genome according to *S. cerevisiae* ORF coordinates to account for three reciprocal translocations between *S. cerevisiae* and *S. uvarum*. Chromosomes are plotted in alternating light and dark purple. Red indicates a *S. cerevisiae* copy number variant and blue indicates a *S. uvarum* copy number variant. For specific coordinates of copy number variants, see Table 1. **A.** Evolved hybrid clones in glucose-limitation all show amplification of *S. uvarum HXT6/7*. **B.** Evolved hybrid clones in phosphate-limitation show a variety of copy number variants, including whole chromosome amplification of *S. cerevisiae* chrIV (3/6 clones) and loss of heterozygosity of part of chromosome XIII (3/6 clones). **C.** Evolved hybrid clones in sulfate-limitation all exhibit amplification of a small region containing *SUL1*.

**Fig S4: Loss of heterozygosity in evolved *S. cerevisiae* clones**

Allele frequency as determined from sequencing read depth data is plotted across chromosomes for each observed loss of heterozygosity event in evolved *S. cerevisiae* clones in **A.** glucose-limited conditions, **B.** phosphate-limited conditions, and **C.** sulfate-limited conditions. The ancestral sequence is plotted at the top of each figure in grey, followed by each sequenced clone from that condition.

## SUPPLEMENTARY TABLES

**Supplemental Table 1: Strain list**

**Supplemental Table 2: Sequencing coverage**

**Supplemental Table 3: Primers used**

